# Express Arm Responses Appear Bilaterally on Upper-limb Muscles in an Arm Choice Reaching Task

**DOI:** 10.1101/2021.09.24.461726

**Authors:** Sarah L. Kearsley, Aaron L. Cecala, Rebecca A. Kozak, Brian D. Corneil

## Abstract

When required, humans can generate very short latency reaches towards visual targets, like catching a falling cellphone. During such rapid reaches, express arm responses are the first wave of upper limb muscle recruitment, occurring ~80-100 ms after target appearance. There is accumulating evidence that express arm responses arise from signaling along the tecto-reticulo-spinal tract, but the involvement of the reticulo-spinal tract has not been well-studied. Since the reticulospinal tract projects bilaterally, we studied whether express arm responses would be generated bilaterally. Human participants (n = 14; 7 female) performed visually guided reaches in a modified emerging target paradigm where either arm could intercept the target. We recorded electromyographic activity bilaterally from the pectoralis major muscle. Our analysis focused on target locations where participants reached with the right arm on some trials, and the left arm on others. In support of the involvement of the reticulospinal tract, express arm responses persisted bilaterally regardless of which arm reached to the target. The latency and magnitude of the express arm response did not depend on whether the arm was chosen to reach or not. However, on the reaching arm, the magnitude of the express arm response was correlated to the level of anticipatory activity. The bilateral generation of express arm responses supports the involvement of the reticulo-spinal tract. We surmise that the correlation between anticipatory activity and the magnitude of express arm responses on the reaching arm arises from convergence of cortically-derived signals with a parallel subcortical pathway mediating the express arm response.

**New and Noteworthy:** Express arm responses have been proposed to arise from the tecto-reticulo-spinal tract originating within the superior colliculus, but the involvement of the reticulo-spinal tract has not been well studied. Here, we show these responses appear bilaterally in a task where either arm can reach to a newly appearing stimulus. Our results suggest that the most rapid visuomotor transformations for reaching are performed by a subcortical pathway.

## Introduction

When time is of the essence, like when catching a cellphone knocked off a desk, visuomotor transformations can occur at times approaching the minimal afferent and efferent conduction delays. A useful marker for these rapid visuomotor transformations is the express arm response. The express arm response, which has also been termed the stimulus locked response (1) or rapid visual response (2), is a burst of upper-limb muscle recruitment that consistently occurs ~100ms after stimulus appearance, regardless of the reach reaction time (RT) (1, 3, 4). The term express arm response was coined to reflect the shared properties of this aspect of upper-limb muscle recruitment with the visual burst of visuomotor neurons in the intermediate and deep layers of the superior colliculus, and with express saccades (5). Express saccades, express arm responses, and the visual burst of visuomotor neurons are all directed toward the location of a visual stimulus, regardless of instructions to move in the opposite direction (4, 6–8). All three responses are also preferentially evoked by stimuli composed of low spatial frequencies and high contrast (9–12). Further, the magnitudes of both express arm responses and the visual burst of visuomotor neurons are inversely related to the ensuing RT (1, 4, 6, 13). These shared properties support the hypothesis that express arm responses are mediated by the superior colliculus (1, 4, 9, 12).

In non-human primates (14), the communication between the superior colliculus and spinal cord is likely indirect, with an interface in the reticular formation. Consistent with this potential interface, express arm responses in humans are augmented by non-visual stimuli thought to excite the reticular formation (2). A distinctive feature of the reticular formation is its extensive, almost equal, bilateral projections to upper-limb muscles (15–17). Cortical motor areas also project bilaterally, but the proportion of ipsilateral cortico-spinal motor projections is lower (18, 19), and muscle responses evoked from ipsilateral motor areas tend to have longer latencies and smaller magnitudes (20–22). To date, express arm responses have been studied only in unimanual reaching tasks. The goal of this study is to test whether express arm responses would be expressed bilaterally when either arm can be used to reach to a visual target.

Previous work has shown an emerging target paradigm, wherein a moving target transiently disappears and then emerges from behind a barrier, elicits robust express arm responses in the reaching arm in almost every participant (5, 12, 23, 24). Here, we modified this paradigm by increasing the number of potential locations of target emergence and allowing the subject to reach toward the emerging target with either arm. These modifications elicited reaches by either the left or right arm for different target locations, and at certain locations elicited left arm reaches on some trials and right arm reaches on other trials. Muscle recruitment for reaches toward these latter locations is critical for our primary aim, which is to determine whether the expression of express arm responses depended on whether the arm was chosen to reach to the target or not. Further, as our task requires participants to choose which arm to move toward the emerging target, a secondary aim was to determine when limb muscle activity indicated whether the associated arm would reach to the target or not. In doing so, we can assess the presence or absence of any relationship between the commitment to move a particular arm and the express arm response. Overall, we found that express arm responses evolved on both the chosen and non-chosen arm. We also found that the time at which limb muscle recruitment indicated which arm would reach to the target was highly variable and was unrelated to the timing of express arm responses. These findings are consistent with express arm responses being relayed through the reticular formation along a tecto-reticulo-spinal pathway and illustrate a surprising degree of independence between the expression of express arm responses and the decision to commit to moving one arm or the other.

## Methods and Materials

### Participants

15 participants (8 males, 7 females; mean age: 21.8 years SD: 1.9) provided informed written consent, were paid for their participation, and were free to withdraw from the experiment at any time. All participants had normal or corrected-to-normal vision, with no current visual, neurological, or musculoskeletal disorders. All participants completed the short form Edinburgh Handedness Inventory (25, 26) which indicated 12 participants were right-handed, 2 mixed-handed, and 1 left-handed. All procedures were approved by the Health Science Research Ethics Board at the University of Western Ontario. One participant (left-handed male) was excluded due to a failure to follow task instruction, as they routinely initiated arm movements before target emergence.

### Apparatus

Participants generated reaching movements with their left and right arms in a bimanual KINARM end-point robot (BKIN Technologies, Kingston, ON, Canada). Movements were generated in the horizontal plane via two handles through shoulder and elbow flexion and extension. A custom built-in projector (ProPixx projector, VPixx, Saint-Bruno, QC, Canada) generated visual stimuli onto an upward facing mirror, located at approximately shoulder height. All visual stimuli were white (110 cd/m^2^) presented against a black (.6 cd/m^2^) background (contrast ratio: 183:1). A shield below the mirror occluded direct vision of the hands, but real-time hand positions were represented via two white dots each with a diameter of 1.5 cm (which equates to approximately 1 degree of visual angle). Throughout the experiment, constant forces of 2 N towards the participant and 5N outward for each hand were applied to increase tonic activity in the pectoralis major (PEC) muscle.

### Experimental Design

Participants completed a modified version of the emerging target paradigm (23) (**Figure 1A**). Participants initiated each trial by bringing the white dots representing their left and right hands into a round, 2cm diameter white starting position, located 45 cm in front of them, and 23 cm to the left and right of center respectively. These starting positions disappeared once the trial was initiated. Although eye movements were not measured, participants were instructed to fixate on a notch in the barrier, 47 cm in front of them, until the target re-emerged. Simultaneous with the start of the trial, a white target (1.5 cm diameter) located above an occluder began moving toward the participant at 15 cm/s. The target disappeared behind the occluder for a fixed duration of 1.5 s before emerging in motion at 15 cm/s below the occluder at one of 7 locations, appearing either at the horizontal center of the occluder, or 3, 7, or 17 cm to the left or right of this central position. Target motion was vertical both before and after disappearance behind the occluder, regardless of where the target emerged. Thus, the time between target disappearance and appearance was fixed at 1.5 s for all target locations. The target was only presented in its entirely after it moved beneath the occluder, preventing the presentation of a half-moon stimulus with a lower overall area. Upon target emergence, participants were instructed to reach toward the emerging target as quickly as possible and were told that they could use either arm to do so. Participants completed four blocks of 350 trials each, with each block containing 50 pseudorandomly intermixed repetitions of each location, yielding a total of 200 trials for each target location.

**Figure 1.**
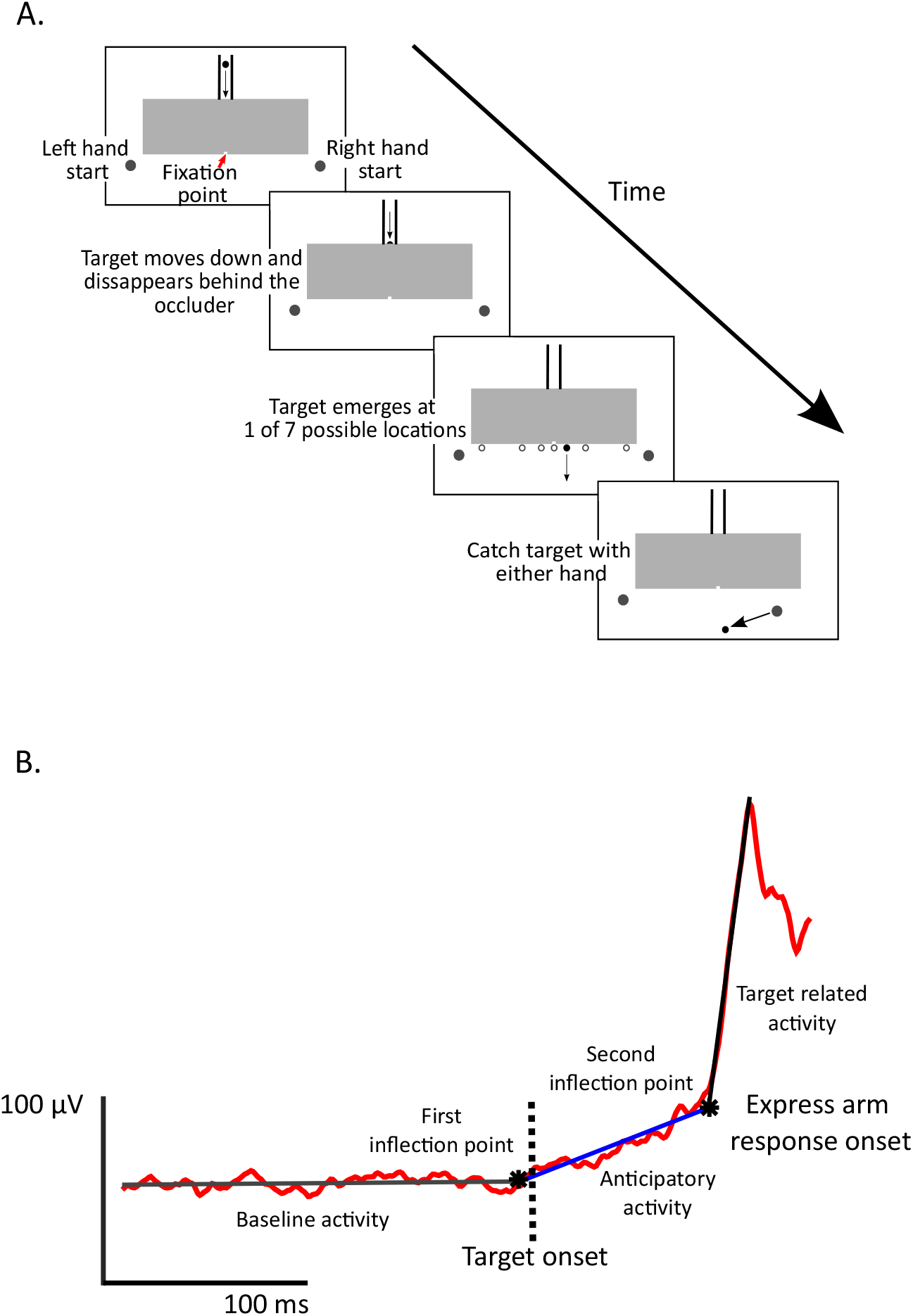
Modified emerging target paradigm and method for classifying express arm responses. A) At the start of each trial, the target appears above an occluder (grey box), and the participant brings their right and left hands into the start position. Then while the participant fixates on a notch in the occluder, the target then moves down the chute, disappears briefly behind the occluder, and then re-emerges below the occluder at one of seven different locations (possible target locations are shown, but these were not presented to the subject). Participants reached to intercept the target with either the right or left arm. B) For this example the muscle activity was fit with a three-piece linear regression, differentiating recruitment during a baseline, anticipatory, and target-related interval. In this case, the time of the second inflection between anticipatory and target-related activity represents the start of the express arm response onset.

### Data acquisition and analysis

Surface electromyographic (EMG) activity was recorded from the clavicular head of the right and left pectoralis major muscle (PEC) with double-differential surface electrodes (Delsys Inc. Bagnoli-8 system, Boston, MA, USA). To ensure consistency, the same individual placed electrodes on the right and left PEC for all participants, using anatomical landmarking and muscle palpation to determine location. EMG signals were amplified by 1000, sampled by the KINARM data system at 1000 Hz, then full wave rectified off-line. Kinematic data was also sampled at 1000 Hz by the KINARM data system. At the time of target emergence, a visual stimulus unseen by the subject was also presented to a photodiode, and all EMG and kinematic data were aligned to this time.

To allow cross-muscle comparisons, we normalized EMG activity to baseline, dividing EMG activity on each trial by the average EMG activity between −500 to −100ms before target onset across all trials. Normalized muscle activity was only used when comparing the magnitudes of recruitment across different muscles, otherwise, source EMG voltages were analyzed.

RT was calculated as the time from target appearance below the occluder, indicated by the photodiode, to the initiation of the reaching movement by the arm that intercepted the target. The reach RT for each trial was determined using a custom MATLAB (version 2014b, The MathWorks, Inc., Natick, Massachusetts, United States) script that found the time when the hand exceeded 5% of its peak velocity of the hand after target onset, and then moved backwards in time to find the point at which hand acceleration following target onset exceeded the 95% confidence interval of acceleration data taken from a period of 100 ms before to 50 ms after target onset. The offset of hand motion was the time at which hand velocity fell below 5% of its peak velocity. The onset and offset of movements were confirmed offline by an analyst in a graphical user interface and adjusted if necessary. We excluded trials with RTs less than 100 ms due to presumed anticipation, and trials with RTs exceeding 500 ms due to presumed inattentiveness. 16% of trials were excluded using these RT constraints, primarily due to anticipatory movements. We also excluded trials consisting of multiple movement segments toward the target, excluding ~2% of trials.

Arm-choice was defined simply as the arm that intercepted the target. A psychometric function was generated using the proportion of right arm reaches as a function of target location. For each participant a logistic regression was fit to the data, using the ‘logit’ MATLAB function: f(p) = log(p/(1-p)), where p is the proportion of right arm reaches. Using the fitted curve, we estimated the theoretical point where a target would be intercepted with either the left or right arm with equal likelihood. The closest target location to this point, referred to as the target of subjective equality, was then used for further analyses, as this target location permitted the best within-muscle comparison of recruitment when that arm was chosen to reach to the target or not.

Previous work examining the express arm response has used a time-series receiver-operating characteristic analysis, contrasting EMG activity for movements into or away from a muscle’s preferred direction (1, 27). Because a given arm only moved in one direction in our study (e.g., all targets lay to the left or right of the right or left arm, respectively), we developed a novel method for detecting and quantifying the express arm response. Our method involves a two- or three-piece linear regression, fitting lines to EMG activity in a baseline, anticipatory (only used for the three-piece linear regression), and post-target interval (see (5, 13) for methods based on a two-piece linear fit). Our rationale for using a three-piece linear regression was based on qualitative observation of mean EMG recruitment, which often started to increase in an anticipatory fashion above baseline before and just after target appearance (**Figure 1B**).

To determine the presence or absence of an express arm response, we took the following steps. First, we ensured that there were at least 25 reaches from a given arm to a particular target (only one arm was used to reach to most target locations). Whenever there were enough reaches from a given arm, we further analyzed the muscle activity from both the left and right PEC, as this provides us with EMG activity from both the reaching and non-reaching arm. We then fit the mean EMG activity spanning from 200 ms before target onset to the time of the peak EMG activity within 125 ms after target onset with a two-piece linear regression. This involved finding the inflection point that minimized the sum of error squares (the loss), delineating baseline activity (spanning from −200 ms to the first inflection point), and the target-related interval (from the inflection point to the peak EMG activity). This two-piece regression sufficed for situations where there was no increasing anticipatory activity between baseline and target related activity. To account for situations where anticipatory activity was present, we fit the data with a three-piece linear regression, enforcing a minimum of 10 ms between the first and third pieces. Doing so involved finding two inflections points that minimized the loss, delineating the baseline activity (spanning from −200 ms to the first inflection point), anticipatory activity (spanning from the first to second inflection point), and the target-related interval (spanning from the second inflection point to the peak EMG activity; see **Figure 1B**). As a three-piece linear regression always decreases loss compared to a two-piece linear regression, we determined whether a three-piece regression would be warranted by calculating the ratio of the loss between the two- and three-piece linear regressions. If the ratio was below 0.7, we used the three-piece linear regression. If the ratio was above 0.7, we used the two-piece linear regression. We also calculated the loss ratio between the two-piece linear regression and regular linear regression. A two-piece linear regression was used if the loss ratio was below 0.6, otherwise a linear regression was used.

Following these steps, we then determined the presence of an express arm response in the following manner. First, the EMG data had to be fit by either a two- or three-piece linear regression; EMG data fit by a linear regression signified the absence of an express arm response. Second, the target related inflection point had to occur within 70-105 ms, and the slope of the first and second piece for a two-piece linear regression, or the second and third piece for a three-piece linear regression had to be significantly different at P < 0.05, as determined by a bootstrapping procedure. If these criteria were met, the latency of the express arm response was defined as the time of the inflection point for the two-piece linear regression, or the second inflection point for the three-piece linear regression. The express arm response magnitude was defined as the difference of the peak EMG activity over the next 15ms to the EMG activity at the onset of the response. We also quantified muscle activity immediately preceding the express arm response (in the results, we term this the “anticipatory activity” for simplicity, although we recognize that anticipatory and baseline activity are equivalent for a two-piece linear regression). Anticipatory activity was quantified as the difference between the EMG signal immediately before the express arm response, and the baseline activity.

In a separate analysis to determine at what point muscle activity reflected arm choice, we used a time-series receiver-operating characteristic (ROC) analysis from EMG activity recorded when participants reach to the target of subjective equality. This target location provided a large sample of EMG activity from a given muscle on trials where the associated arm or the opposite arm reached to the target. We separated EMG activity based on which arm reached to the target, then analyzed at every time sample (1 ms) from 500ms before target onset to the end of the trial. For each time-point we calculated the area under the ROC curve, which is the probability that an ideal observer could discriminate whether the associated arm would reach to the target or not, based solely on the EMG activity. Values of 1 or 0 indicate perfectly correct or incorrect discrimination respectively, whereas a value of 0.5 indicates chance discrimination. We set the threshold discrimination at 0.6 because this criterion exceeded the 95% confidence intervals determined previously using a bootstrapping procedure (13). The time of discrimination was defined as the first point in time at which the ROC value exceeded 0.6 for at least eight of ten subsequent time-samples.

### Statistical Analysis

Statistical analyses were performed in MATLAB. To compare the proportion of participants generating an express arm response (termed express arm response prevalence) as a function of muscle, arm choice, and location, a chi-squared test was used, and Bonferroni corrected when necessary. A paired t-test was used to compare the latency and magnitude of the express arm response within a muscle at the target of subjective equality. We relied on non-normalized EMG for our magnitude analysis for within muscle comparisons, and EMG activity normalized to baseline for across muscle comparisons. For all correlational or unpaired t-test analyses, one value per participant was included in the analysis. In situations where there was more than one observation for a given participant the response of the participant was taken as the average of all observations, as suggested in (28).

## Results

The reticular formation is a likely interface in a tectal pathway mediating express arm responses. Given the bilateral projections from the reticular formation, we wondered whether express arm responses would be expressed bilaterally in a task where participants could choose which arm to use to intercept an emerging target. We recorded muscle activity from the right and left PEC muscles as participants completed a modified emerging target paradigm (**Figure 1A**). Targets could emerge at one of seven locations below the barrier, and participants reached to catch the target as fast as possible with either arm. We analysed muscle activity from both the reaching and non-reaching arm to determine the presence of the express arm response. We also examined the time at which muscle activity indicated that the associated arm would reach toward the target or not, relative to the time of any express arm response.

### Arm-choice as a function of target location, and defining the target of subjective equality

Participants were free to choose which arm to move for all targets but tended to choose the arm closest to the emerging target (**Figure 2)**. We quantified participant behaviour by fitting a psychometric curve to the proportion of right arm reaches expressed as a function of target location. The point of subjective equality defines the theoretical target location where a participant would reach with one arm on half of all trials, and with the other arm on the other half of trials. From the point of subjective equality, we found the closest actual target location, referred to as the target of subjective equality, for each participant (see **Figure 2A** for a representative subject). This location was associated with a high number of reaches from either arm in all participants. Across our sample, the target of subjective equality was at center (n = 10), 3 cm left (n = 2) or 7 cm left (n = 2) of center (**Figure 2B**). The target of subjective equality permits a within-muscle comparison of recruitment when the associated arm was chosen to reach or not. In general, locations other than the target of subjective equality did not generate enough reaches from both arms for within muscle comparisons.

**Figure 2.**
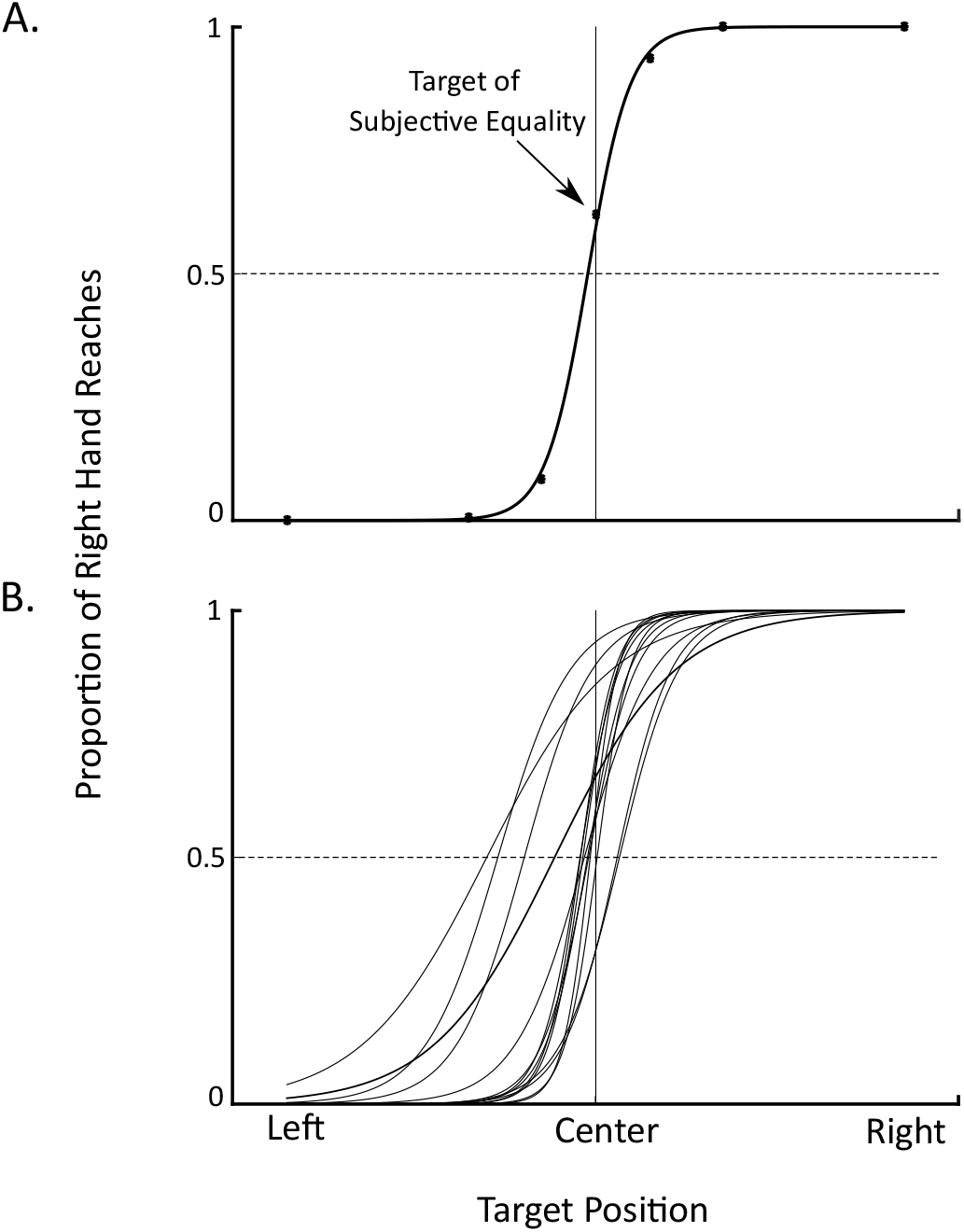
Arm Choice as a Function of Target location. A: A single participant example of right arm choice as a function of target location. Each black dot represents a location where the target emerged on a subset of the trials. A psychometric function was fit to the data and the target of subjective equality was chosen as the target closest to the horizontal dash line. B: Psychometric functions for all participants.

### Do express arm responses appear bilaterally?

The main question we wanted to address was whether express arm responses evolve bilaterally when either arm could be used to intercept an emerging target. **Figure 3A** shows the average muscle activity from an exemplar participant (same participant as **Figure 2A**), across all positions where at least 25 reaches were made by the associated arm. These data show how participants tended to reach with the arm closest to the target (e.g., note how the right or left arm tended to reach for targets in the right or left hemifield, respectively). Using either a two-piece or three-piece linear regression to determine whether there was an express arm response (**Figure 1B**, see Methods), we observed express arm responses in both the reaching and non-reaching arm (inflection points are denoted by the black dot; express arm responses in **Figure 3A** are denoted by the first or second dots when a two- or three-piece linear regression was used, respectively). When detected, express arm responses occurred ~90ms after target appearance in both the reaching and non-reaching arms.

**Figure 3.**
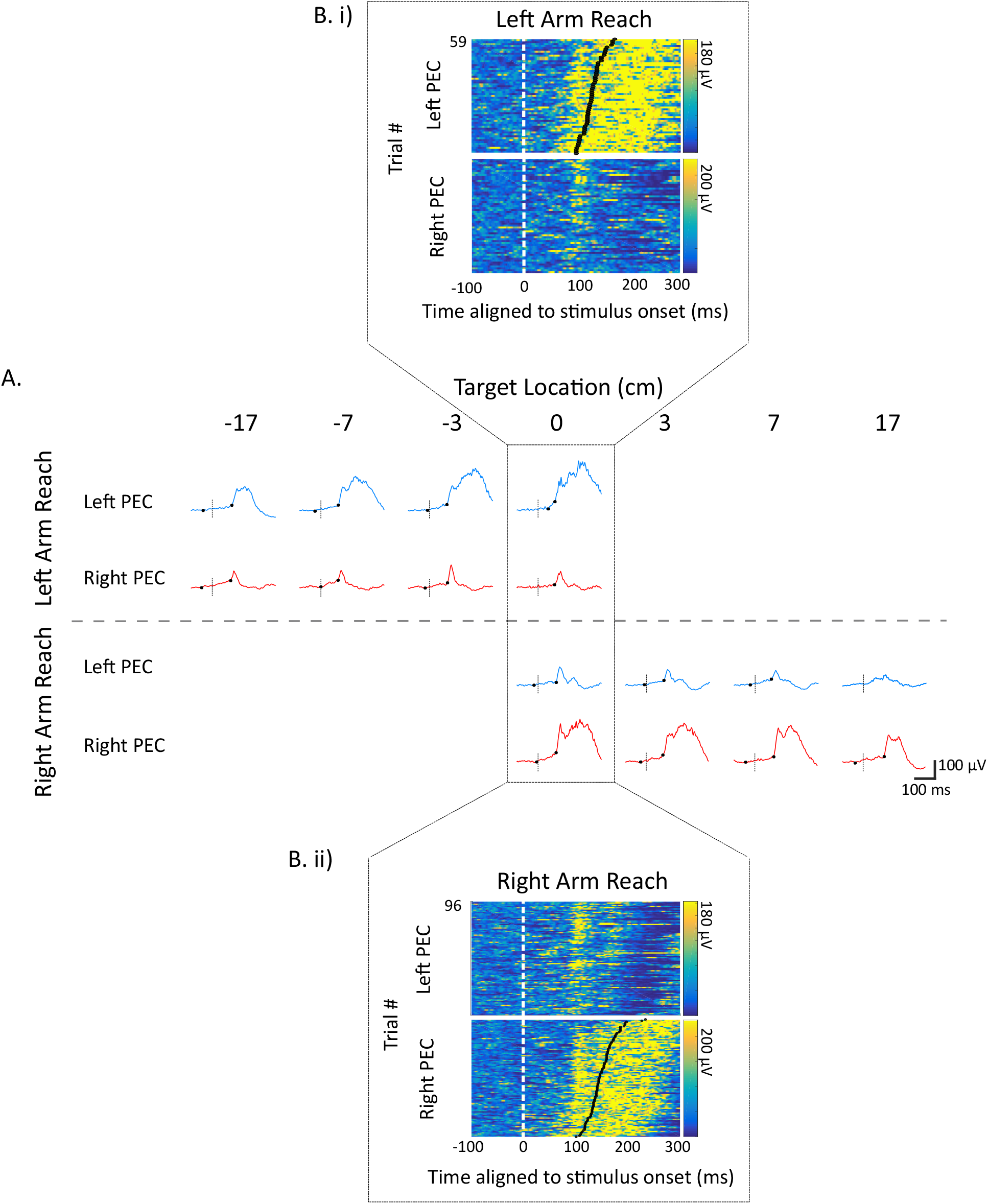
Bilateral muscle recruitment in a representative participant. A) Average muscle activity (+/- SE) for all reaches as a function of target location. Averages are plotted only if there were at least 25 trials where the given arm reached to the target. The 0 cm location is the target of subjective equality, as this featured many trials where either the right or left arm reached to the target. Stimulus onset indicated by the black vertical dotted line. Black dots represent the inflection points, the first or second of which indicate the time at which an express arm response was detected when using the two- or three-piece linear regression respectively (see methods for further details). B) Depiction of trial-by-trial recruitment from left (top) and right (bottom)

Previous reports have emphasized that the trial-by-trial timing of express arm responses is more aligned to stimulus rather than movement onset (1, 4). As shown in **Figure 3B,** we indeed found that the timing of express arm responses was more tied to stimulus rather than movement onset, regardless of whether the associated arm reached or not. This characteristic feature of express arm responses appears as the vertical banding of EMG activity in **Figure 3B** when muscle activity is aligned to stimulus onset, showing a burst of muscle recruitment ~90 ms after target emergence regardless of the ensuing reach RT. Following this bilateral generation of the express arm response, a more prolonged period of increased recruitment was observed only on the muscle associated with the reaching arm.

The prevalence of express arm responses is known to vary across paradigms and participants (1, 5, 9, 12). We wanted to know whether all participants had express arm responses in general, and further whether the responses were equally prevalent in the reaching and non-reaching arms. As shown in **Figure 4**, the modified emerging target paradigm elicited express arm responses from at least one participant at each location. Further, almost all participants (n = 13) generated express arm responses following target presentation to at least one location. Compared to the null-hypothesis that the response only occurs in the reaching arm, we found that the response also occurred in the non-reaching arm (Chi-squared test: p < 0.001, c2= 52.3858, df=1). We also compared the prevalence of express arm responses in the reaching and non-reaching arm grouped across all locations, and further at each location individually. Using a chi-squared test we found express arm responses occurred more frequently in the reaching arm compared to the non-reaching arms across all locations (p= 0.002, c2= 9.6671, df=1). Thus, although express arm responses can evolve bilaterally on both upper limbs, they are more likely to occur in the reaching arm. Comparing express arm responses at each location revealed they were less likely to occur at the 17 cm locations (Chi-squared test, Bonferroni corrected, alpha = 0.0083, p < 0.0083). No other differences were found based on location.

**Figure 4.**
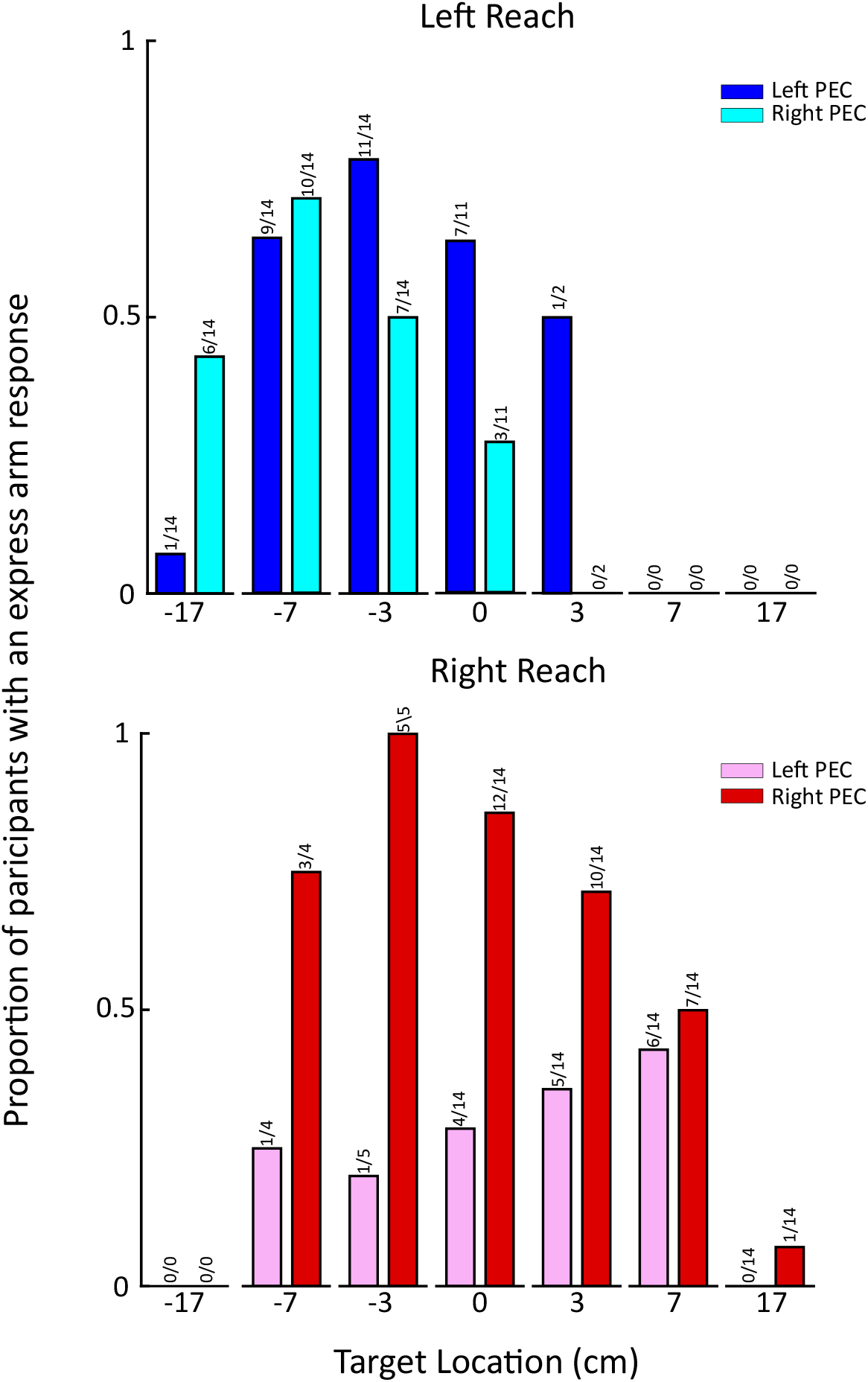
Proportion of subjects exhibiting an express arm response as a function of arm and target location. At each target location prevalence is determined as the proportion of participants exhibiting an express arm response relative to the number of subjects who generated enough reaches with the given arm at that particular location (recall at least 25 reaches had to be made by a given arm for the analysis of the express arm response).

### Properties of express arm responses

Next, we were interested in the latency and magnitude of express arm responses recorded bilaterally, and whether these measures differed depending on whether the associated arm was selected to move or not. Previous work has shown that express arm response latency (9) and/or magnitude (4) may differ depending on stimulus properties and task context. We therefore examined these properties at the target of subjective equality, as a function of whether the associated arm was chosen to reach or not (note that this is a within-muscle comparison). Using only paired observations (i.e., when express arm responses were detected in a given muscle regardless of whether the arm was chosen to move or not) we found no difference in express arm response latency with arm choice (paired observations shown as the connected points in **Figure 5A;** p = 0.5299, t = −0.6565, df = 8). Further, using a single factor ANOVA we found no difference in response latency across target locations (p > 0.05). These results reinforce the qualitative observation from **Figure 3A** that the express arm response evolves consistently ~90 ms irrespective of arm choice. Along with latency, magnitude was also not significantly different when the arm was chosen or not chosen to reach at the point of equal selection (**Figure 5B;** p = 0.1485, t = 1.5989, df = 8), or across target locations (single factor ANOVA, p > 0.05).

**Figure 5.**
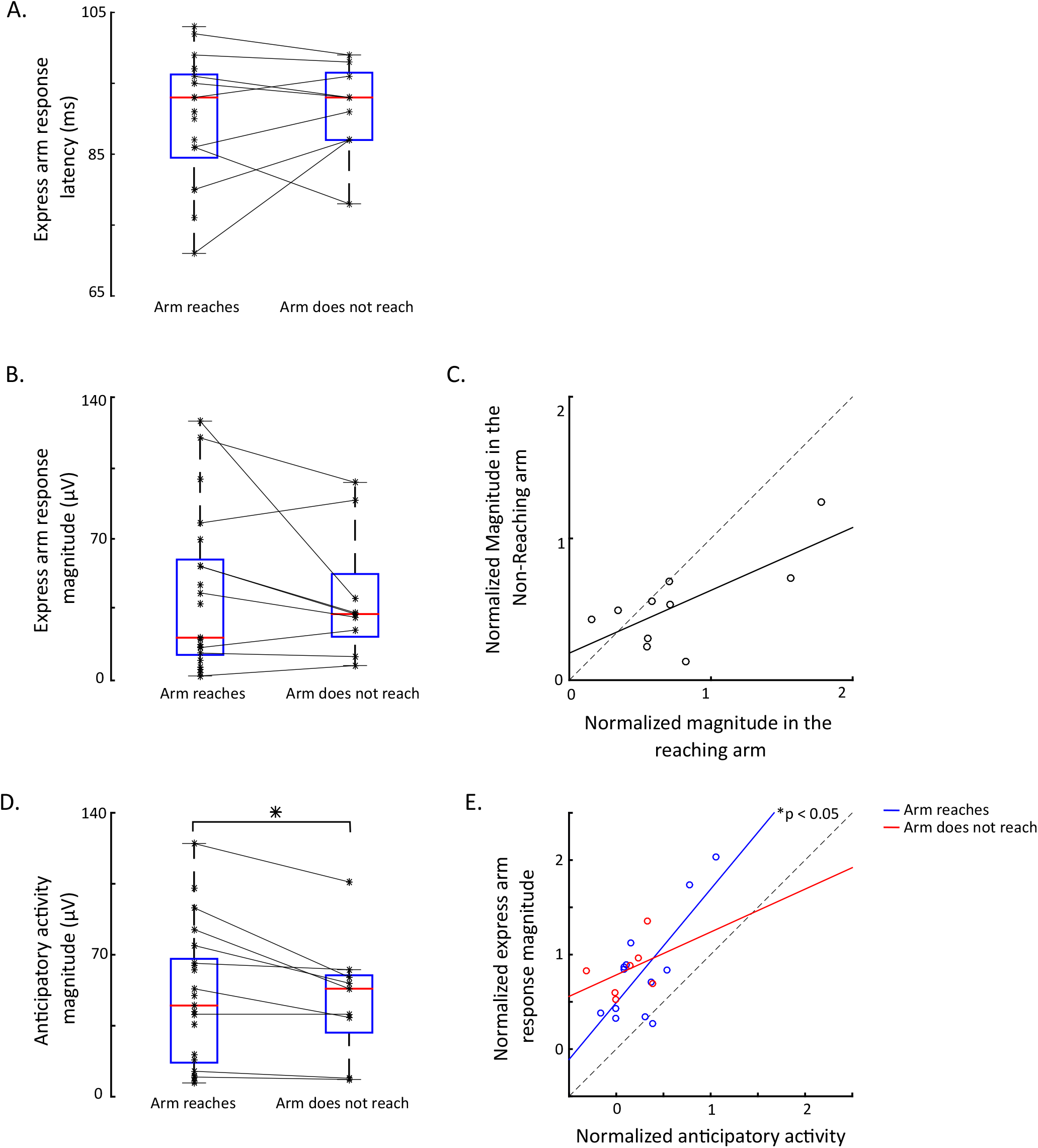
Analyses of the characteristics of express arm response. The latency (A) or magnitude (B) of the express arm response as a function of whether the associated arm reached or not, taken from the target of equal selection. Lines connect within-muscle observations, points not connected by lines show data which did not have a paired value and were not included in the analyses. C) The magnitude of the express arm response in the reaching and non-reaching arm are significantly correlated across participants (r = 0.7141, p = 0.0204). Each dot represents a unique combination of target and subject where express arm responses were observed on both the reaching and non-reaching arm. The black line indicates the linear regression fit, and the dashed line represents the line of unity. D) Anticipatory activity, measured as the level of EMG activity just prior to the express arm response. Same format as A. Anticipatory activity was significantly higher when the arm was selected to reach to the target (p = 0.0080). E) Correlation of the level of anticipatory activity to the magnitude of the express arm response for the reaching (blue, r = 0.7602, p = 0.0026) and non-reaching (red, r = 0.3777, p > 0.05) arms. Each dot represents an observation, with the black line indicating the linear regression fit.

We also investigated whether the express arm response on the reaching arm was different as a function of whether the non-reaching arm also showed an express arm response. To test this, we compared the latency and magnitude of the express arm response in the reaching arm when the non-reaching arm also exhibited an express arm response versus when the express arm response was only observed in the reaching arm. Using a student’s t-test, we found no difference in the express arm response latency or magnitude on the reaching arm as a function of the presence or absence of an express arm responses on the non-reaching arm (latency: p > 0.05, t = 0.4947, df = 6.4457, magnitude: p > 0.05, t =1.5089, df = 6.9517; data not shown).

If mediated by a common source like the reticular formation, we would expect the magnitude of express arm responses on the reaching and non-reaching arm to be correlated across participants and targets (e.g., a larger express arm response on the reaching arm should be associated with a larger express arm response on the non-reaching arm). To analyze this, we identified target locations where an express arm response was observed on both the reaching and non-reaching arm, using one observation for each participant (see Methods), and found that express arm response magnitudes were positively correlated between the muscles (**Figure 5C,** Pearson correlation, p = 0.0204, r = 0.7141; every point represents a unique observation for a participant; note magnitudes are normalized here since this is an across-muscle comparison). Thus, larger express arm response magnitudes on the reaching arm tended to be associated with larger express arm response magnitudes on the non-reaching arm. Interestingly, on average, the magnitude of the express arm responses was about 1.5 times as large on the reaching versus non-reaching arm (p = 0.0631, t = 2.1190, df = 9).

In our paradigm, participants knew in advance that targets would appear medial relative to the starting position of both the left and right arm, leading us to wonder if participants anticipated which arm to use prior to target emergence. We analyzed the potential influence of such anticipation and found greater anticipatory activity when the associated arm was chosen to reach to the target of subjective equality (**Figure 5D**; paired t-test, p = 0.0080, t = 3.5082, df = 8). This relationship between anticipatory activity and arm choice can be seen in **Figure 3A** on the right PEC at the 0 cm target; note how anticipatory activity preceding the express arm response was greater when the right rather than left arm reached to the target. This level of anticipatory activity related to the magnitude of the ensuing express arm response in the reaching arm (n.b., the latter measure quantifies the EMG magnitude above anticipation; **Figure 5E** blue points; r = 0.7602, p = 0.0026). However, this relationship was not seen in the non-reaching arm (**Figure 5E,** red points; r = 0.3777, p > 0.05), potentially due to the smaller sample size. Thus, the level of anticipatory activity attained just before the express arm response related to the magnitude of the express arm response on the reaching arm.

### When, relative to the express arm response, does muscle activity relate to arm choice?

The preceding analyses showed that greater levels of anticipatory muscle recruitment relate to the choice to use the associated arm to reach to the target. These results lead us to wonder when muscle activity predicts which arm was going to move, and whether this time relates in a systematic way to the latency or expression of an express arm response. To address this, we performed a time-series ROC analysis to compare the muscle activity when the arm was chosen to reach or not and searched for the time at which an ideal observer could correctly discriminate arm choice from such EMG activity (see Methods). **Figure 6A** shows one example of this analysis, showing the average activity of left PEC muscle for the exemplar participant (same participant as **Figure 2A** and **Figure 3**) preceding left or right arm reaches to the target of subjective equality (top plot, blue or pink traces respectively), as well as the associated time-series ROC (bottom plot). For this example, the discrimination time at which EMG activity reliably predicted which arm would reach was 69 ms after target onset, which preceded the express arm response. Across our entire sample, and regardless of whether participants exhibited an express arm response or not, we observed no systematic relationship between the discrimination time indicating which arm would move and the latency of express arm responses, with discrimination times variably preceding, occurring within, or following the express arm response epoch (**Figure 6B**). We also observed no obvious relationship between this discrimination time and the generation of express arm responses; subjects exhibited express arm responses regardless of whether the discrimination time occurred earlier or later than the express arm response. This analysis reveals a lack of any relationship between aspects of muscle recruitment reflecting arm choice and expression of the express arm response.

**Figure 6.**
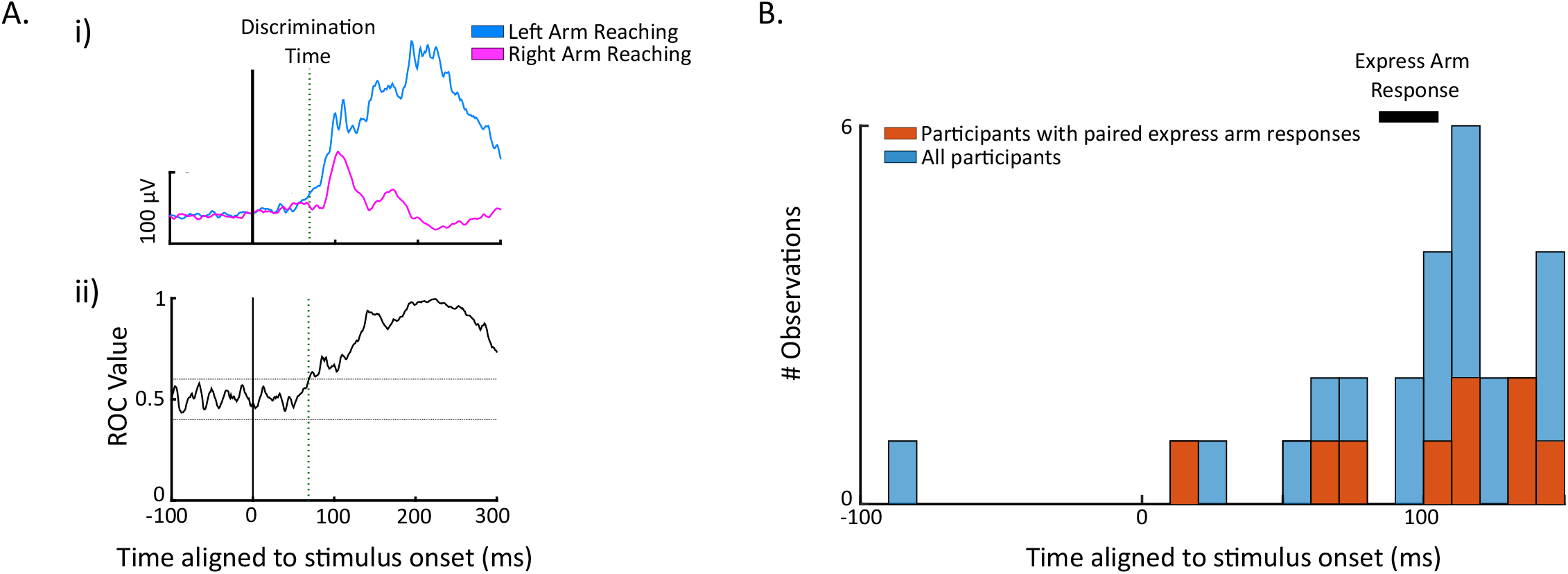
Time of arm choice discrimination based on muscle activity. A) data from the exemplar participant, with the top plot (i) depicting mean EMG (+/- SE) from left PEC for reaches using the left (blue) or right (pink) arm, and the bottom plot (ii) showing the time-series ROC analysis used to determine the time at muscle activity predicts arm choice. Green vertical dotted lines in the inset represents the time of discrimination (69 ms). B) Histogram of the discrimination times organized into bins of 10ms. Orange bins depict observations where the participant exhibited an express arm response on a given muscle when the associated arm was selected to reach or not. Blue bins depict observations where express arm responses were not observed.

### Kinematic Consequences of the Express arm response

The express arm response is a brief period of muscle recruitment that increases muscle force. Previous work with unimanual anti-reach, delay, or stop-signal tasks has shown that express arm responses can produce small, task inappropriate, movements toward a target (4, 29, 30). The non-reaching arm provides a further opportunity to study the kinematic consequences of express arm responses in isolation from ensuing reach-related activity. First, we looked at the velocity of both the reaching and non-reaching arm at every location and consistently saw a small movement towards the target in the non-reaching arm. This can be seen in **Figure 7A** where we have plotted horizontal velocity from the exemplar participant for both the reaching and non-reaching arms at every location. As expected, the velocity is much higher in the reaching arm than in the non-reaching arm, but there is clearly a small deviation of the non-reaching arm toward the target (represented at an increased scale in the insets in **Figure 7A**). To quantify the non-reaching arm’s peak velocity and allow cross-participant comparisons, we normalized it by the peak velocity of the reaching arm. We found on average the non-reaching arm had a peak velocity that was 8.11 ± 2.27% of the reaching arm. Compared to a null hypothesis that no movement occurs in the non-reaching arm, the non-reaching arm did indeed move towards the stimulus (Student’s t-test, p < 0.001, t = −13.3950, df = 13). Next, we compared the peak velocity in the non-reaching arm based on whether an express arm response was observed but did not find any difference in peak velocity based on whether an express arm response was observed (peak velocity: 8.94 ± 2.22%) or not (peak velocity: 7.80 ± 2.68%) (**Figure 7B;** student’s t-test, p > 0.05). Thus, although the non-reaching arm did move toward the target, the peak velocity of this movement was unrelated to the detection of an express arm response. This is a somewhat surprising result; however, this could be due to a failure of the surface EMGs to reliably detect all express arm responses, especially in situations with a low signal to noise ratio.

**Figure 7.**
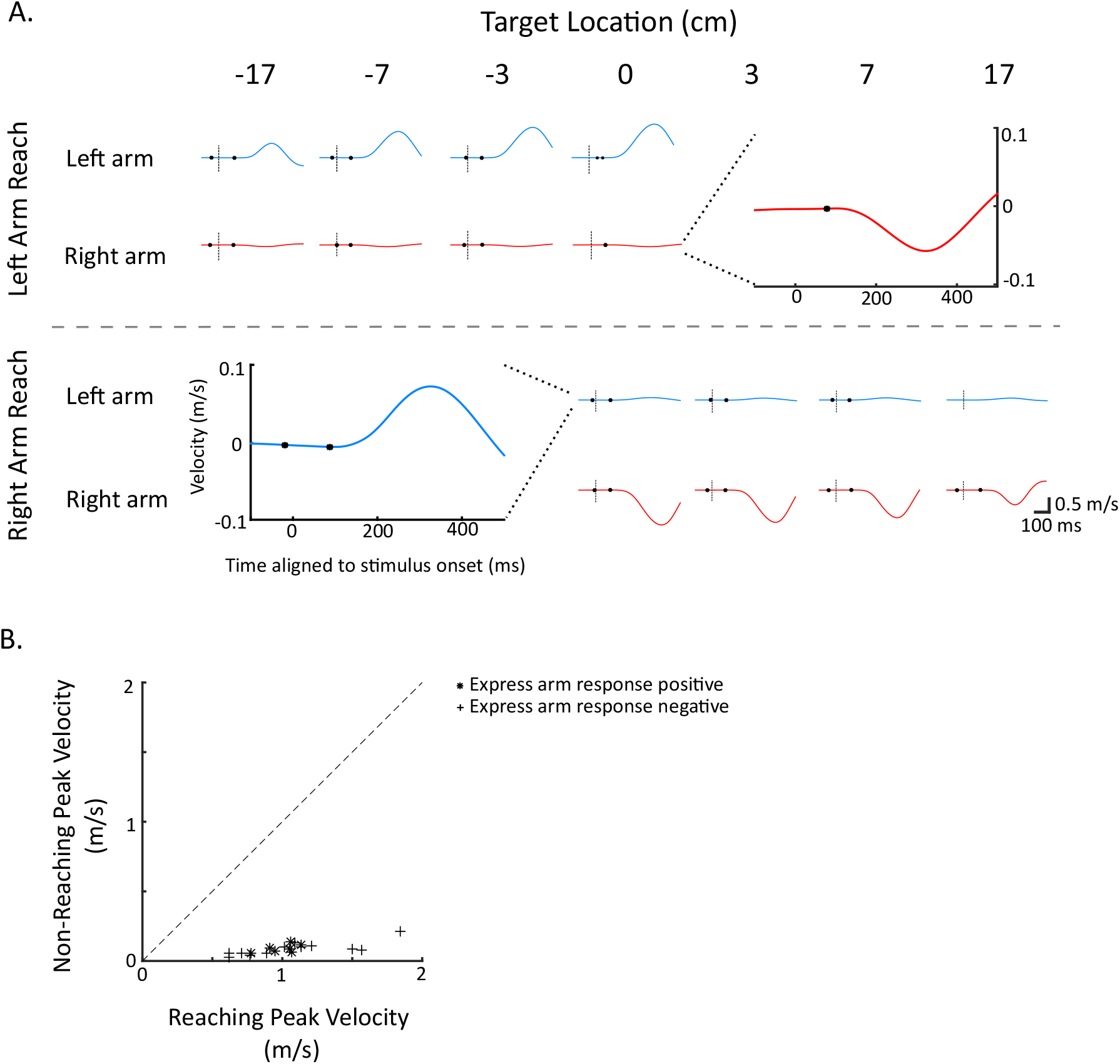
Velocity traces for the exemplar participant. A) Average velocity (+/- SE) for both the reaching and non-reaching arm across locations, with the first or second black dot representing the latency of the express arm response when present when the two- or three-piece linear regression was used to detect the response respectively. Expanded graphs represent the velocity trace from the non-reaching arm at the target of subjective equality, at an enlarged y-axis scale. B) Scatter plot showing the peak velocity of the reaching vs non-reaching arm. Black dashed line shows line of unity and symbols depict whether an express arm response was observed on the non-reaching arm or not.

Another feature that is apparent in the velocity traces of the non-reaching arm is that the small movement toward the target is followed by a brief reversal in velocity. This reversal reflects a small returning movement of the non-reaching arm back toward the starting position. Interestingly, the EMG correlates of this returning movement on the non-reaching arm are apparent in **Figure 3A,** where recruitment levels after the express arm response drop below the levels of anticipatory recruitment attained just before the express arm response.

Given the presence of anticipatory EMG activity, we examined whether the reaching arm drifted slowly inwards, given that all targets appeared medial relative to starting hand positions. To do this, we compared the position of the hand at baseline versus immediately before the express arm response and observed no relationship between the level of anticipatory activity and any change in hand position (p > 0.05). This suggests the anticipatory activity did not move the hand, perhaps because any arising forces were insufficient to overcome the inertia of the hand, or because of co-contraction of unrecorded antagonist muscles.

A key behavioural correlation seen in previous research using unimanual tasks is that larger express arm responses tend to precede shorter-latency RTs (1, 4). Given that this study is the first to study express arm responses in a bimanual task, we examined our data for the presence of any relationships between express arm responses and RTs. We first confirmed that the express arm response magnitude in the reaching arm is negatively correlated to reach RT (left panel of **Figure 8A** shows trial-by-trial data for the right PEC from the exemplar participant; right panel of **Figure 8A** shows that the r-values across all participants with an express response at the target of equal selection lay significantly below zero; average r = −0.3710, p < 0.001, t = 10.7281, df = 12). Next, we examined whether the magnitude of the express arm response on the non-reaching arm related to the RT of the reaching arm, as a common drive mechanism predicts that a larger express arm muscle response on the non-reaching arm should precede shorter latency RTs on the reach arm. However, we found no relationship between the magnitude of the express arm response on the non-reaching arm and the RT of the reaching arm either in the exemplar participant (left panel of **Figure 8B**) or across the sample (the distribution of r-values in right panel in **Figure 8B** does not differ from zero, average r = −0.0016, p >> 0.05, t = 0.036, df = 6). Instead, as we were able to occasionally extract a RT from the movement of the non-reaching arm, we found a weaker negative correlation that approaches significance between non-reaching express arm response magnitude and non-reaching movement RT (left panel of **Figure 8C** for exemplar participant; right panel of **Figure 8C** for the sample; average r = −0.1879, p = 0.0591, t = 2.3246, df = 6). This final negative correlation does suggest a relationship between the express arm response on the non-reaching arm and the RT for the small movement of that arm, even when the other arm intercepts the target.

**Figure 8.**
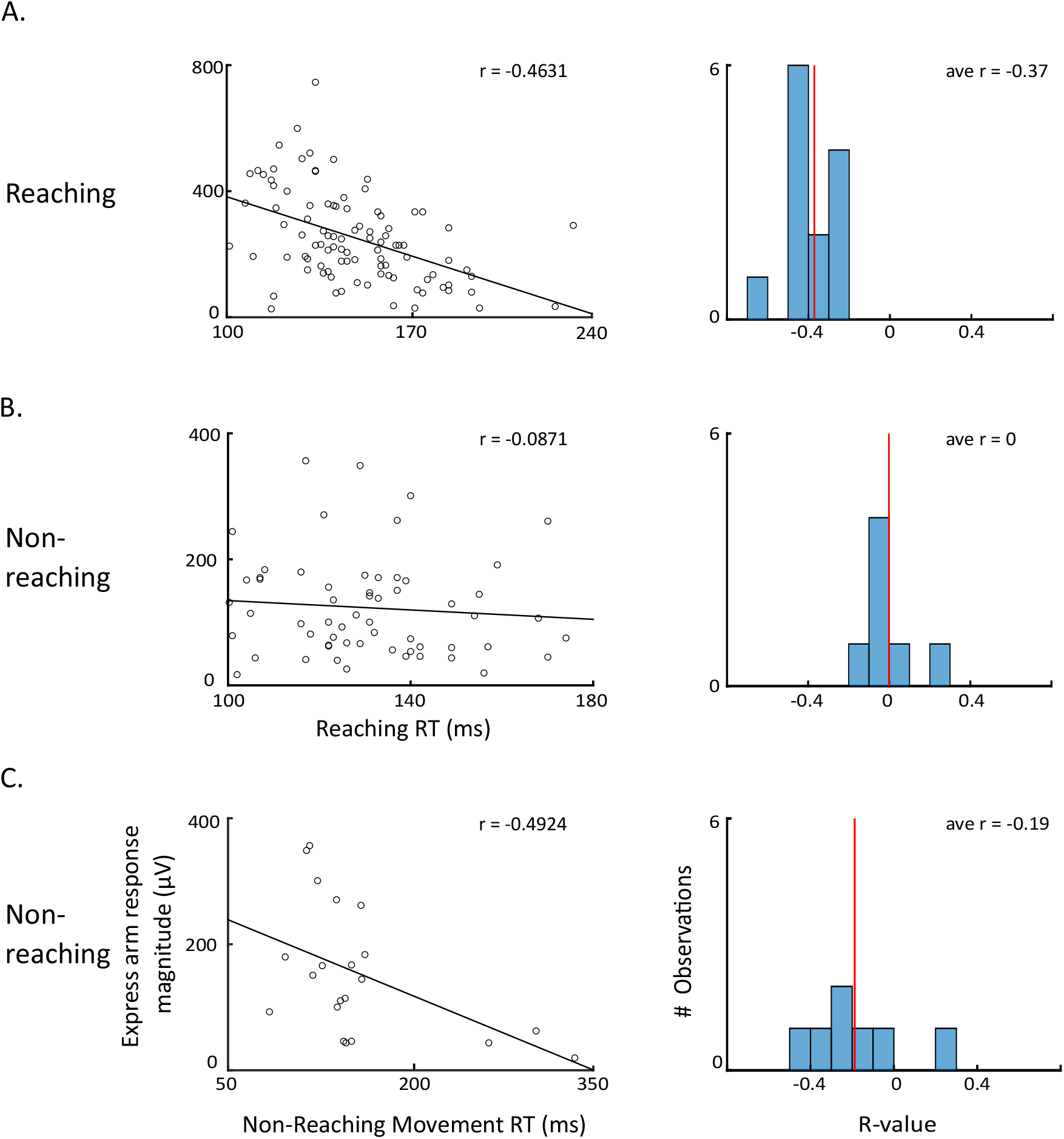
Correlations for express arm response magnitude and reaction time (RT). A) In both the exemplar participant (Left; each point represents data from a single trial) and population (Right) there is a negative trial-by-trial correlation between the magnitude of the express arm response in the reaching arm and the RT of the movement. B) No such negative relationship was observed between the magnitude of the express arm response on the non-reaching arm and the RT of the reaching arm for either the exemplar participant or across the sample. C) A weaker negative correlation was observed between the express arm response on the non-reaching arm and the RT of the non-reaching arm (when a movement was present).

## Discussion

We investigated whether express arm responses occur bilaterally in a task where either arm can be used to intercept a target. We were interested in: i) the prevalence, timing, and magnitude of any express arm responses whether the arm reaches or not, ii) how these measures related to anticipatory muscle recruitment, and iii) how these measures related to the kinematics of any associated movement. Express arm responses occurred on both the reaching and non-reaching arms, and response magnitude interacted with the preceding level of anticipatory activity. Our results are consistent with a reticular interface for signals arising soon after target onset in the superior colliculus, and the interaction of such signals with pre-existing activity related to the anticipation of target emergence.

### Interactions between anticipatory recruitment, the express arm response, and voluntary reach-related activity

In our task, all targets emerged medial to the starting position of each hand. Participants often anticipated target emergence to a degree that influenced muscle recruitment. Such anticipatory recruitment, which we presume has a cortical origin as participants become familiar with the task, influenced the magnitude but not timing of the express arm response; participants with more anticipatory recruitment tended to have larger express arm responses (**Figure 5E)**, and anticipatory recruitment tended to be larger on the arm chosen to reach (**Figure 5D**). The relationships between anticipatory recruitment and express arm responses resemble gain scaling seen for the spinal stretch reflex following a mechanical perturbation of the arm (31). Recruitment from subsequent longer-loop reflexes may not be gain-scaled if it were to be counterproductive to the task at hand. A future line of research should investigate whether the express arm response indeed exhibits gain scaling. This could be done by varying the loading force on the muscle of interest and investigating how anticipatory recruitment influences both the express arm response and ensuing phases of recruitment.

The time-series ROC analysis shown in **Figure 7** shows greater anticipatory activity in some participants on the arm that reached to the target, showing a degree of commitment before target presentation. Such anticipatory recruitment is only a bias, as participants still reached with hand closest to the most peripheral targets (**Figure 2**). This bias may result from trial history or fatigue (e.g., favor one arm if the other arm was used on the previous trial). A bias favoring one arm may explain the lack of a relationship between the magnitude of the express arm response on the non-reaching arm and the RT of the reaching arm (**Figure 8B**), as a common drive to both muscles would predict a negative relationship between the express arm response magnitude of either arm and the reach RT. Instead, since the magnitude of the express arm response is also influenced by anticipatory activity, a bias in anticipatory activity against the non-reaching arm may have muted the magnitude of the ensuing express arm response.

Previous work shows that larger express arm responses precede shorter RTs (1, 4). We observed this on the reaching arm (**Figure 8A**), and this relationship on the non-reaching arm (when a reaction time for the non-reaching arm could be extracted) approached significance (**Figure 8C**). Comparing the evolution of muscle activity on the reaching versus non-reaching arm is quite interesting; whereas express arm responses are readily apparent on both, subsequent phases of more prolonged recruitment are only observed on the reaching arm. The kinematics of movement of the non-reaching arm provides an opportunity to better understand the kinetic consequences of the relatively brief express arm response, and like previous results (4, 29, 30), the express arm response is associated with a small movement of the non-reaching arm toward the target followed by a reversal in the voluntary response epoch. This reaffirms that, despite the relatively brief nature of the express arm response, it imparts a kinetic consequence.

### The express arm response: a low-level reflex potentiated by high-level processes

Circumstantial evidence suggests that express arm responses arise from signalling along a tecto-reticulo-spinal pathway, initiated from the intermediate or deep layers of the superior colliculus (1, 4, 9, 16, 29). The related phenomenon of express neck muscles responses (13, 27, 32) have been directly correlated to visual responses in the intermediate and deep superior colliculus (33). Many of the key response properties of express arm responses resemble those of express saccades, in which the role of the superior colliculus is well understood (34, 35). Express saccades are a low-level reflex that are, somewhat paradoxically, potentiated by high-level processes (6, 36). Cortical inputs raise collicular activity in advance of the visual response, and express saccades occur when the sum of such pre-target activity and the visual response exceed saccade threshold (37). There is no one cortical area solely responsible for such pre-target activity, but instead inputs from many frontal and parietal areas are thought to converge on the superior colliculus (38–41). If similar mechanisms govern express arm responses in our paradigm, then cortical inputs related to implied motion and the timing of target appearance would increase the pre-target activity within the superior colliculus. Express arm responses have been potentiated in other behavioural paradigms that presumably engage different top-down inputs into the superior colliculus (3, 23, 24, 29).

This perspective on express arm responses blurs the distinctions between concepts such as target selection, movement planning or preparation, execution, and the commitment to reach with a given arm. In the emerging target task, which engages a high degree of preparation and anticipation, we speculate that EMG activity arises from converging inputs from multiple descending pathways, each of which has distinct characteristics and dynamics during different phases of a trial. Anticipatory EMG activity is characterized by recruitment that can be biased in favour of one arm or the other, starts before target presentation, and is not directed to a particular target. The express arm response is driven by and directed toward a particular target (and hence is arguably synonymous with both target selection and execution) and can be distributed bilaterally. Finally, the commitment to reach with a particular arm is completed after the express arm response and consists of unilateral recruitment of one arm and simultaneous relaxation of the other. Fundamentally, the networks governing muscle recruitment in the context of this task are likely nested (42–44), and as the output of the pathway that most rapidly links vision to action, the express arm response offers a unique window where target-related muscle activity is solely driven by subcortical descending pathways.

### Neural substrates for the express arm response

The interface between the superior colliculus and motor periphery is likely indirect, and our work here adds to a small body of literature that more has considered the potential involvement of other interfaces. For example, Glover and Baker (2019)(2) reported enhanced express arm responses (what they termed rapid visual responses) in a unimanual response task when visual stimuli were combined with other auditory, vestibular, or somatosensory stimuli. Such non-visual stimuli are thought to enhance responses in the reticular formation, hence they attributed the facilitation they observed on express arm responses to the influence of such non-visual stimuli in the reticular formation. Very rapid on-line corrections can also be shared across the upper and lower limbs, presumably via subcortical pathways (45). Further, by combining transcranial brain stimulation and electrical stimulation of the median nerve, Nakajima, Suzuki and colleagues (46, 47) proposed that rapid limb responses to changing visual inputs arose from integration within cervical interneurons of corticospinal inputs with visual information rapidly relayed along a subcortical tectoreticulospinal pathways. Whether cervical interneurons are involved in the generation of express arm responses, perhaps in conjunction with the reticular formation, remains to be determined but this seems likely given the broad convergence between descending motor pathways (42).

How visual information reaches the intermediate and deep layers of the superior colliculus is unclear. Visual information could be relayed directly to the superficial superior colliculus along retinotectal projections, and then access deeper layers via intracollicular pathways (48). Visual signals could also access the intermediate and deep layers via projections from striate and extrastriate cortices (49), including areas such as MT (48, 50). Regardless, any route conveying visual information to the intermediate and deep layers of the superior colliculus must do so very rapidly.

But are there alternative pathways for express arm responses? Most reports of time- and direction-locked visual responses in monkey motor cortex have latencies ~100 ms (51, 52), which lag express arm responses. However, there are reports of rapid visual responses ~50-60 ms in the primate motor cortex (53, 54), so theoretically visual signals may have rapid enough access to the motor cortex. But could rapid visual responses in motor cortex then evoke bilateral and simultaneous recruitment of upper arm musculature? While ipsilateral corticospinal projections tend to be relatively sparse (18, 19) and slower conducting (20–22, 55), the motor cortex could theoretically receive ipsilateral and contralateral visual information from MT(56–58). Recent work has also shown strong, bilateral projections from motor cortex to the reticular formation (59). While a corticospinal or corticoreticulospinal route for express arm responses is theoretically possible, visual responses in motor cortex (unlike those in the superior colliculus) have not been studied in sufficient detail to enable comparison with the known properties of express arm responses.

### Comparison to past studies and methodological considerations

Our study is the first to investigate express arm responses when either arm could reach toward a target. Further, we increased the number of potential targets from two to seven. Despite these changes, all but one participant exhibited an express arm response to at least one target. We attribute this to our paradigm maintaining implied motion behind the barrier and a high degree of certainty about the time of target emergence, which have been suggested to be the main factors increasing express arm response prevalence and magnitude (5, 12, 23).

Participants chose which arm reached to the target, doing so as quickly as possible. Previous work has shown that arm choice tends to reflect the hemifield of target appearance, with a slight bias to use the dominant hand for central targets (60, 61). Past versions of a hand-choice task did not instruct participants to reach as fast as possible (60, 61), thus the dominant hand could have been used for all targets. Instead, hand choice still largely reflected the hemifield of presentation.

Our task found, for each subject, a target location eliciting reaches with the right arm on some trials, and the left arm on others. Doing so enabled comparison of muscle activity as a function of whether the associated arm was selected to reach or not for movements to the same visual target. For most participants (n = 10), this target of equal selection was the center target. Assuming participants followed task instruction, this center target would be ~1 degree below the fovea. While foveal visual stimuli are represented bilaterally in the superior colliculus (62), a variety of reasons make it unlikely that this could explain our observations of bilateral express arm responses. First, bilateral responses were obtained for the four participants who had off-centre targets of equal selection (two participants at each of 3 or 7 cm to the left, equating to ~3 or 7 degrees); such visual targets are represented unilaterally in the superior colliculus. Second, targets beside the target of equal selection still provoked bilateral responses; it was simply that reaches to these locations were predominantly done by one arm. Third, past work dissociating initial eye and hand position have shown that the express arm responses encode the location of the visual stimulus relative to the current position of the hand, not the eye (3).

In our paradigm, the retinal image of the central target moved more rapidly than more peripheral targets. However, we did not find any influence of target location on the magnitude of express arm responses on either the reaching or non-reaching arm. Previous work has reported that faster moving targets evoke larger express arm responses (12), but the range of retinal velocities used in our experiment may not have been large enough to reveal this effect. Related work by Cross and colleagues requiring on-line corrections following a jump in cursor position also found that the earliest visuomotor responses are invariant for jumps that are greater than 2 cm in magnitude (63). The lack of any relationship between target location and express arm response magnitude is therefore not surprising, although future work should more systematically investigate this question.

In past work, potential targets were positioned to either side of the hand, and express arm responses were detected via time-series ROC analyses of the increases or decreases in muscle activity following target presentation into or out of the muscle’s preferred direction of movement. Here, all targets lay medial to the starting position of the hand, in the preferred direction for pectoralis major. We therefore developed a new method for detecting express arm responses, relying on two- or three-piece linear regressions fit to mean EMG activity (see METHODS). This method was not perfect, and we had the impression of false negatives, particularly when trying to detect smaller express arm responses on the non-reaching arm and when recordings tended to be noisier. When express arm responses were detected, EMG activity displayed the characteristic trial-by-trial changes more aligned to target rather than movement onset (e.g., **Figure 3B**).

Our positioning of targets medial to both hands, with loading forces in the opposite direction, meant that pectoralis major was the only muscle on which the bilateral distribution of express muscle responses could have been assessed. Having established this, future experiments should record other limb muscles, and require contraction of a given muscle in one arm and relaxation of the same muscle on the other arm to reach to a target (e.g., by altering loading forces or changing initial posture). Indeed, the most common bilateral recruitment profile evoked by stimulation of the reticular formation is ipsilateral muscle facilitation and contralateral muscle suppression (64). If the pathway mediating the bilateral distribution of express muscle responses is to have any functional benefit, it should be able to flexibly map target locations onto different combinations of bilateral muscle recruitment.

Finally, our participants were either right-handed (n = 12) or ambidextrous (n =2). Previous studies of express arm responses studied few left-handed participants (1, 5, 12), but there has been no suggestion of differences between left- and right-handed participants. We speculate that the express arm response would remain bilateral in left-hand dominant participants, but this remains to be determined.

### Conclusions

Our work contributes to the understanding of express arm responses, showing for the first time to our knowledge that the underlying pathway distributes the motor signal bilaterally. Our results are consistent with the reticular formation serving as an interface between the superior colliculus and the motor periphery. Our overall hypothesis is that signalling along the tectoreticulospinal pathway initiates the first wave of limb muscle recruitment in circumstances requiring rapid visually-guided reaching. We are mindful of the convergence of cortical inputs into all nodes of this pathway, including the superior colliculus, the reticular formation, spinal interneuron networks, and the motoneuron. Rather than being directly involved in express arm responses, cortical inputs into these subcortical nodes, for example with anticipatory signals that bias arm choice, may dampen or augment the vigor of the earliest visually-related responses. Further characterization of express arm responses, and the integration of such signalling with task-relevant information, can more precisely constrain the neural mechanisms for express arm responses, and address the integration of such signalling with cortical inputs to initiate and guide our most rapid visually-guided behaviours.

## Acknowledgements

This work is supported by Discovery Grants to BDC from the Natural Sciences and Engineering Research Council of Canada (NSERC; RGPIN 311680 and 04394-2021) and an Operating Grant to BDC from the Canadian Institutes of Health Research (CIHR; MOP-93796). SLK was supported by a NSERC CGS-M. RAK was supported by an Ontario Graduate Scholarship. The equipment used in this experiment was purchased using funds from the Canadian Foundation for Innovation. Additional support came from the Canada First Research Excellence Fund (BrainsCAN). We thank Drs. Tim Carroll, Gerald Loeb, and Andrew Pruszynski, as well as Sam Contemori, Rechu Divakar and Dr. Yang Chao, for helpful feedback on earlier versions of the manuscript.

## References

1. Andrew Pruszynski J, King GL, Boisse L, Scott SH, Randall Flanagan J, Munoz DP. Stimulus-locked responses on human arm muscles reveal a rapid neural pathway linking visual input to arm motor output. European Journal of Neuroscience 32: 1049–1057, 2010. doi: 10.1111/j.1460-9568.2010.07380.x.

2. Glover IS, Baker SN. Multimodal stimuli modulate rapid visual responses during reaching. Journal of Neurophysiology 122: 1894–1908, 2019. doi: 10.1152/jn.00158.2019.

3. Gu C, Pruszynski J, Gribble P, Corneil B. Done in 100 ms: path-dependent visuomotor transformation in the human upper limb. Journal of neurophysiology 119: 1319–1328, 2018. doi: 10.1152/JN.00839.2017.

4. Gu C, Wood DK, Gribble PL, Corneil BD. A trial-by-trial window into sensorimotor transformations in the human motor periphery. Journal of Neuroscience 36: 8273–8282, 2016. doi: 10.1523/JNEUROSCI.0899-16.2016.

5. Contemori S, Loeb GE, Corneil BD, Wallis G, Carroll TJ. The influence of temporal predictability on express visuomotor responses. Journal of Neurophysiology 125: 731–747, 2021. doi: 10.1152/JN.00521.2020.

6. Pare M, Munoz DP. Saccadic reaction time in the monkey: advanced preparation of oculomotor programs is primarily responsible for express saccade occurrence. Journal of Neurophysiology 76: 3666–3681, 1996. doi: 10.1152/JN.1996.76.6.3666.

7. Fischer B, Ramsperger E. Human express saccades: extremely short reaction times of goal directed eye movements. Experimental Brain Research 57: 191–195, 1984.

8. Everling S, Dorris MC, Klein RM, Munoz DP. Role of Primate Superior Colliculus in Preparation and Execution of Anti-Saccades and Pro-Saccades. Journal of Neuroscience 19: 2740–2754, 1999. doi: 10.1523/JNEUROSCI.19-07-02740.1999.

9. Kozak RA, Kreyenmeier P, Gu C, Johnston K, Corneil BD. Stimulus-locked responses on human upper limb muscles and corrective reaches are preferentially evoked by low spatial frequencies. eNeuro 6, 2019. doi: 10.1523/ENEURO.0301-19.2019.

10. Ludwig CJH, Gilchrist ID, McSorley E. The influence of spatial frequency and contrast on saccade latencies. Vision Research 44: 2597–2604, 2004. doi: 10.1016/J.VISRES.2004.05.022.

11. Chen C-Y, Sonnenberg L, Weller S, Witschel T, Hafed ZM. Spatial frequency sensitivity in macaque midbrain. Nature Communications 2018 9:1 9: 1–13, 2018. doi: 10.1038/s41467-018-05302-5.

12. Kozak RA, Corneil BD. High-contrast, moving targets in an emerging target paradigm promote fast visuomotor responses during visually guided reaching. Journal of Neurophysiology 126: 68–81, 2021. doi: https://doi.org/10.1152/jn.00057.2021.

13. Goonetilleke SC, Katz L, Wood DK, Gu C, Huk AC, Corneil BD. Cross-species comparison of anticipatory and stimulus-driven neck muscle activity well before saccadic gaze shifts in humans and nonhuman primates. Journal of Neurophysiology 114: 902–913, 2015. doi: 10.1152/jn.00230.2015.

14. Nudo RJ, Masterton RB. Descending pathways to the spinal cord: II. Quantitative study of the tectospinal tract in 23 mammals. Journal of Comparative Neurology 286: 96–119, 1989. doi: 10.1002/cne.902860107.

15. Grantyn A, Grantyn R. Axonal Patterns and Sites of Termination of Cat Superior Colliculus Neurons Projecting in the Tecto-Bulbo-Spinal Tract. Experimental Brain Research 46: 243–256, 1982.

16. Corneil BD, Munoz DP. Overt responses during covert orienting. Neuron 82 Cell Press: 1230–1243, 2014.

17. Davidson AG, Schieber MH, Buford JA. Bilateral spike-triggered average effects in arm and shoulder muscles from the monkey pontomedullary reticular formation. Journal of Neuroscience 27: 8053–8058, 2007. doi: 10.1523/JNEUROSCI.0040-07.2007.

18. Rosenzweig ES, Brock JH, Culbertson MD, Lu P, Moseanko R, Edgerton VR, Havton LA, Tuszynski MH. Extensive spinal decussation and bilateral termination of cervical corticospinal projections in rhesus monkeys. The Journal of comparative neurology 513: 151–163, 2009. doi: 10.1002/CNE.21940.

19. Armand J, Kuypers HGJM. Cells of origin of crossed and uncrossed corticospinal fibers in the cat: a quantitative horseradish peroxidase study. Experimental brain research 40: 23–34, 1980. doi: 10.1007/BF00236659.

20. Montgomery LR, Herbert WJ, Buford JA. Recruitment of ipsilateral and contralateral upper limb muscles following stimulation of the cortical motor areas in the monkey. Experimental Brain Research 230: 153–164, 2013. doi: 10.1007/s00221-013-3639-5.

21. Ziemann U, Ishii K, Borgheresi A, Yaseen Z, Battaglia F, Hallett M, Cincotta M, Wassermann EM. Dissociation of the pathways mediating ipsilateral and contralateral motor-evoked potentials in human hand and arm muscles. The Journal of Physiology 518: 895–906, 1999. doi: 10.1111/J.1469-7793.1999.0895P.X.

22. MacKinnon CD, Quartarone A, Rothwell JC. Inter-hemispheric asymmetry of ipsilateral corticofugal projections to proximal muscles in humans. Experimental Brain Research 157: 225–233, 2004. doi: 10.1007/s00221-004-1836-y.

23. Kozak R, Corneil B. Stimulus-locked responses on human upper limb muscles prefer low spatial frequency, high contrast, and fast moving targets. Journal of Vision 20: 554, 2020. doi: 10.1167/jov.20.11.554.

24. Contemori S, Loeb GE, Corneil BD, Wallis G, Carroll TJ. Trial-by-trial modulation of express visuomotor responses induced by symbolic or barely detectable cues. Journal of neurophysiology 126: 1507–1523, 2021. doi: 10.1152/JN.00053.2021.

25. Oldfield RC. The Assessment and Analysis of Handedness: The Edinburgh Inventory. Neuropsychologia 9: 97–113, 1971.

26. Veale JF. Edinburgh Handedness Inventory - Short Form: A revised version based on confirmatory factor analysis. Laterality 19: 164–177, 2014. doi: 10.1080/1357650X.2013.783045.

27. Corneil BD, Olivier E, Munoz DP. Visual responses on neck muscles reveal selective gating that prevents express saccades. Neuron 42: 831–841, 2004. doi: 10.1016/S0896-6273(04)00267-3.

28. Makin TR, de Xivry JJO. Ten common statistical mistakes to watch out for when writing or reviewing a manuscript. eLife 8, 2019. doi: 10.7554/ELIFE.48175.

29. Wood DK, Gu C, Corneil BD, Gribble PL, Goodale MA. Transient visual responses reset the phase of low-frequency oscillations in the skeletomotor periphery. European Journal of Neuroscience 42: 1919–1932, 2015. doi: 10.1111/EJN.12976.

30. Atsma J, Maij F, Gu C, Medendorp WP, Corneil BD. Active Braking of Whole-Arm Reaching Movements Provides Single-Trial Neuromuscular Measures of Movement Cancellation. Journal of Neuroscience 38: 4367–4382, 2018. doi: 10.1523/JNEUROSCI.1745-17.2018.

31. Pruszynski JA, Kurtzer I, Lillicrap TP, Scott SH. Temporal Evolution of “Automatic Gain-Scaling.” Journal of Neurophysiology 102: 992–1003, 2009. doi: 10.1152/JN.00085.2009.

32. Corneil BD, Munoz DP, Chapman BB, Admans T, Cushing SL. Neuromuscular consequences of reflexive covert orienting..

33. Rezvani S, Corneil B. Recruitment of a head-turning synergy by low-frequency activity in the primate superior colliculus. Journal of neurophysiology 100: 397–411, 2008. doi: 10.1152/JN.90223.2008.

34. Edelman J, Keller E. Activity of visuomotor burst neurons in the superior colliculus accompanying express saccades. Journal of neurophysiology 76: 908–926, 1996. doi: 10.1152/JN.1996.76.2.908.

35. Dorris M, Paré M, Munoz D. Neuronal activity in monkey superior colliculus related to the initiation of saccadic eye movements. The Journal of neuroscience : the official journal of the Society for Neuroscience 17: 8566–8579, 1997. doi: 10.1523/JNEUROSCI.17-21-08566.1997.

36. Schiller PH, Haushofer J, Kendall G. How do target predictability and precueing affect the production of express saccades in monkeys? The European journal of neuroscience 19: 1963–1968, 2004. doi: 10.1111/J.1460-9568.2004.03299.X.

37. Munoz DP, Dorris MC, Paré M, Everling S. On your mark, get set: Brainstem circuitry underlying saccadic initiation 1. 2000.

38. Chen M, Liu Y, Wei L, Zhang M. Parietal cortical neuronal activity is selective for express saccades. The Journal of neuroscience : the official journal of the Society for Neuroscience 33: 814–823, 2013. doi: 10.1523/JNEUROSCI.2675-12.2013.

39. Dash S, Peel TR, Lomber SG, Corneil BD. Impairment but not abolishment of express saccades after unilateral or bilateral inactivation of the frontal eye fields. Journal of neurophysiology 123: 1907–1919, 2020. doi: 10.1152/JN.00191.2019.

40. Johnston K, Koval MJ, Lomber SG, Everling S. Macaque dorsolateral prefrontal cortex does not suppress saccade-related activity in the superior colliculus. Cerebral cortex (New York, NY : 1991) 24: 1373–1388, 2014. doi: 10.1093/CERCOR/BHS424.

41. Dash S, Peel TR, Lomber SG, Corneil BD. Frontal eye field inactivation reduces saccade preparation in the superior colliculus but does not alter how preparatory activity relates to saccades of a given latency. eNeuro 5, 2018. doi: 10.1523/ENEURO.0024-18.2018.

42. Alstermark B, Isa T. Circuits for skilled reaching and grasping. Annual review of neuroscience 35: 559–578, 2012. doi: 10.1146/ANNUREV-NEURO-062111-150527.

43. Fautrelle L, Bonnetblanc F. On-line coordination in complex goal-directed movements: A matter of interactions between several loops. Brain Research Bulletin 89: 57–64, 2012. doi: 10.1016/J.BRAINRESBULL.2012.07.005.

44. Reynolds RF, Day BL. Direct visuomotor mapping for fast visually-evoked arm movements. Neuropsychologia 50: 3169–3173, 2012. doi: 10.1016/J.NEUROPSYCHOLOGIA.2012.10.006.

45. Fautrelle L, Prablanc C, Berret B, Ballay Y, Bonnetblanc F. Pointing to double-step visual stimuli from a standing position: very short latency (express) corrections are observed in upper and lower limbs and may not require cortical involvement. Neuroscience 169: 697–705, 2010. doi: 10.1016/J.NEUROSCIENCE.2010.05.014.

46. Nakajima T, Ohtsuka H, Irie S, Suzuki S, Ariyasu R, Komiyama T, Ohki Y. Visual information increases the indirect corticospinal excitation via cervical interneurons in humans. https://doi.org/101152/jn004252020 125: 828–842, 2021. doi: 10.1152/JN.00425.2020.

47. Suzuki S, Nakajima T, Irie S, Ariyasu R, Ohtsuka H, Komiyama T, Ohki Y. Subcortical Contribution of Corticospinal Transmission during Visually Guided Switching Movements of the Arm. Cerebral Cortex 00: 1–17, 2021. doi: 10.1093/cercor/bhab214.

48. May PJ. The mammalian superior colliculus: laminar structure and connections. Progress in brain research 151: 321–378, 2006. doi: 10.1016/S0079-6123(05)51011-2.

49. Tigges J, Tigges M. Distribution of retinofugal and corticofugal axon terminals in the superior colliculus of squirrel monkey. Investigative Ophthalmology & Visual Science 20: 149–158, 1981.

50. Maunsell JHR, van Essen DC. The connections of the middle temporal visual area (MT) and their relationship to a cortical hierarchy in the macaque monkey. The Journal of neuroscience : the official journal of the Society for Neuroscience 3: 2563–2586, 1983. doi: 10.1523/JNEUROSCI.03-12-02563.1983.

51. Riehle A. Visually induced signal-locked neuronal activity changes in precentral motor areas of the monkey: hierarchical progression of signal processing. Brain research 540: 131–137, 1991. doi: 10.1016/0006-8993(91)90499-L.

52. Shen L, Alexander GE. Neural correlates of a spatial sensory-to-motor transformation in primary motor cortex. Journal of neurophysiology 77: 1171–1194, 1997. doi: 10.1152/JN.1997.77.3.1171.

53. Kilavik BE, Confais J, Ponce-Alvarez A, Diesmann M, Riehle A. Evoked potentials in motor cortical local field potentials reflect task timing and behavioral performance. Journal of neurophysiology 104: 2338–2351, 2010. doi: 10.1152/JN.00250.2010.

54. Reimer J, Hatsopoulos NG. Periodicity and evoked responses in motor cortex. The Journal of neuroscience : the official journal of the Society for Neuroscience 30: 11506–11515, 2010. doi: 10.1523/JNEUROSCI.5947-09.2010.

55. Tazoe T, Perez MA. Selective Activation of Ipsilateral Motor Pathways in Intact Humans. The Journal of Neuroscience 34: 13924–13934, 2014. doi: 10.1523/JNEUROSCI.1648-14.2014.

56. Abe H, Tani T, Mashiko H, Kitamura N, Hayami T, Watanabe S, Sakai K, Suzuki W, Mizukami H, Watakabe A, Yamamori T, Ichinohe N. Axonal projections from the middle temporal area in the common marmoset. Frontiers in Neuroanatomy 12: 89, 2018. doi: 10.3389/FNANA.2018.00089/BIBTEX.

57. Kolster H, Peeters R, Orban GA. The Retinotopic Organization of the Human Middle Temporal Area MT/V5 and Its Cortical Neighbors. The Journal of Neuroscience 30: 9801, 2010. doi: 10.1523/JNEUROSCI.2069-10.2010.

58. Palmer SM, Rosa MGP. Quantitative Analysis of the Corticocortical Projections to the Middle Temporal Area in the Marmoset Monkey: Evolutionary and Functional Implications. Cerebral Cortex 16: 1361–1375, 2006. doi: 10.1093/CERCOR/BHJ078.

59. Fisher KM, Zaaimi B, Edgley SA, Baker SN. Extensive Cortical Convergence to Primate Reticulospinal Pathways. The Journal of neuroscience : the official journal of the Society for Neuroscience 41: 1005–1018, 2021. doi: 10.1523/JNEUROSCI.1379-20.2020.

60. Bryden PJ, Pryde KM, Roy EA. A performance measure of the degree of hand preference. Brain and Cognition 44: 402–414, 2000. doi: 10.1006/brcg.1999.1201.

61. Bryden MP, Singh M, Steenhuis RE, Clarkson KL. A behavioral measure of hand preference as opposed to hand skill. Neuropsychologia 32: 991–999, 1994. doi: 10.1016/0028-3932(94)90048-5.

62. Chen CY, Hoffmann KP, Distler C, Hafed ZM. The Foveal Visual Representation of the Primate Superior Colliculus. Current Biology 29: 2109–2119.e7, 2019. doi: 10.1016/j.cub.2019.05.040.

63. Cross KP, Cluff T, Takei T, Scott SH. Visual feedback processing of the limb involves two distinct phases. Journal of Neuroscience 39: 6751–6765, 2019. doi: 10.1523/JNEUROSCI.3112-18.2019.

64. Davidson AG, Buford JA. Bilateral actions of the reticulospinal tract on arm and shoulder muscles in the monkey: stimulus triggered averaging. Experimental Brain Research 173: 25–39, 2006. doi: 10.1007/s00221-006-0374-1.

